# DiffDock-Glide: a hybrid physics-based and data-driven approach to molecular docking

**DOI:** 10.1101/2025.06.02.657461

**Authors:** Lukas Herron, Jumana Dakka, Kun Yao, Da Shi, Yuqi Zhang, Steven V. Jerome

## Abstract

Recent years have seen a rise in applications of deep learning to problems in the molecular sciences. Among them, the diffusion model DiffDock stands out as a method for docking small molecules into protein binding sites. But DiffDock struggles to compete with conventional docking methods, especially for targets outside its training set. We develop a hybrid model called DiffDock-Glide which addresses some shortcomings of deep learning docking methods: it uses a modified generative process to generate samples within a binding pocket and the confidence model is replaced with Glide’s post-docking minimization pipeline. We evaluate DiffDock-Glide on the Posebusters dataset and show improved sampling of near-native poses, especially for sequences without homologues in the training set. We also evaluate DiffDock-Glide’s performance in virtual screening compounds from the DUD-E dataset against receptor structures generated by AlphaFold2 and report enrichment values that broadly surpass those from traditional Glide.

## I. INTRODUCTION

Structure-based drug design is a paradigm of drug discovery that uses structural information to rationally design small molecules that bind to targets with high specificity and affinity.^1^ Computational approaches have proven indispensable for reducing the vast space of possible molecules to a small number of potentially high-affinity binders.^2–4^ Historically, the most successful computational strategies have been physics-based molecular docking models that explicitly account for energetic^5–8^ and entropic^9^ contributions to the binding free energy.

But, in recent years, alternative approaches rooted in deep learning have emerged, all of which share a core assumption that the relevant chemical interactions are represented in structural and evolutionary data and can be implicitly modeled by a neural network. In a foundational application of deep learning to biology and chemistry, AlphaFold2 (AF2) modeled protein structure to unprecedented accuracy by training an artificial intelligence system on coevolutionary information and the structural contents of the Protein Data Bank (PDB).^10^ Since AF2 there has been a dramatic increase in deep learning methods for predicting structures that arise from molecular interactions^11,12^, including ones specifically focused on molecular docking.^13,14^

However, comparative studies find that, at present, classical docking methods significantly outperform deep learning ones.^15^ Deep learning methods struggle to predict binding poses for receptors from families outside the training data^15^ and fail to model the molecular interactions that stabilize binding with high fidelity^16^, indicating that the amount and diversity of currently available training data is insufficient to model physical chemistry more accurately than first-principles descriptions.

Classical learning theory attributes the performance gap to deep learning models having many more free parameters than are necessary to capture the latent variables. The bias-variance trade-off quantifies an inverse relationship between model complexity and capacity for generalization: the reduction in fitting error (bias) due to introducing more tunable parameters is concomitant with an increased ability to memorize noise (variance) present in the data.^17^ Deep learning models operate in the low-bias high-variance regime, and as a result they are prone to overfit their training data and struggle to generalize to out-of-distribution settings.^18^

Models in the natural sciences are less prone to overfitting: free parameters are kept to a minimum through the inclusion of conceptual prior knowledge, i.e. inductive biases.^19^ In fact, incorporating task-specific inductive biases into deep learning models is a popular approach to improving generalization ability^20,21^, and generative diffusion models are particularly promising for introducing such biases via modifications to the diffusion dynamics. One may, for instance, define equivariant dynamics^22^, restrict the diffusion to a manifold^23,24^, bias the generative dynamics^25^, or represent thermodynamic quantities.^26,27^

In this work, we present a hybrid physics-based and deep learning approach to molecular docking with biophysical and chemical inductive biases. We use DiffDock^13^–a diffusion model–to sample putative binding poses, then score and refine the poses with Glide^7^– a physics-based molecular docking algorithm. We modify the generative dynamics so that the location of the binding site may be specified in a manner that introduces negligible computational overhead and does not require retraining the model.

We benchmark our method, called DiffDock-Glide, on the Posebusters dataset, demonstrating improved generalization to novel sequences and sampling of near-native binding poses compared to DiffDock alone. We also virtual screen compounds from the DUD-E^28^ dataset against AF2-generated receptor structures and obtain enrichment values that broadly surpass those obtained from Glide or DiffDock alone.

## II. METHOD

The hybrid molecular docking model has two stages. The first stage involves generating a set of potential binding poses using the DiffDock model and a modified generative process that optionally restricts generation to a putative binding site. The second stage involves refining and scoring the generated poses using Glide.

### A. Binding site-conditional DiffDock

DiffDock is a diffusion model that predicts binding poses for a small-molecule given its chemical identity and a target protein structure. It is a *blind* docking algorithm – anywhere inside or on the target protein’s surface is a potential binding site. But, in a realistic structure-based drug discovery setting, one often knows the location of the binding site and would like to explore poses therein. To this end, we bias DiffDock’s generative process towards the location of the site using the particle guidance framework.^25^ We will briefly summarize the core operating principles of DiffDock before introducing the biasing mechanism.

DiffDock incorporates internal symmetries of molecular conformers as inductive biases. During conformer generation, perturbations to molecular degrees of freedom are represented as a product space ℙ of groups: 𝕋(3) *× SO*(3) *× SO*(2)^*m*^; 𝕋(3) is the group of 3D-translations; *SO*(3) is 3D-rotations; and *SO*(2)^*m*^ are 2D-rotations of *m* torsion angles. Clearly, the actions of ℙ on a particular conformer will result in new conformers; it is then natural to ask if two arbitrary conformers are connected through an action of ℙ. Indeed, Corso et. al. show that a typical pair of conformers lie on a manifold ℳ defined by actions of ℙ.^13^

DiffDock is a score-based diffusion model^29^–that is, a stochastic differential equation–with dynamics onℳ:

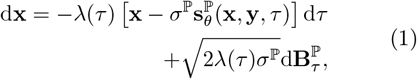

where 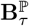 is Brownian motion on ℳ, *λ*(*τ*) is a time dilation factor to accelerate convergence of the diffusion process, and *σ*^ℙ^ is the scale of the noise perturbations. The vectors **x** and **y**–containing the positions of the heavy ligand and *C*_*α*_ protein atoms respectively–are inputs to a score network **s**_*θ*_. The score network acts as a time-dependent vector field on ℳ that generates binding poses from noise vectors by simulating Eq. 1 from *τ* = 0 to *τ* = 1.

The DockGen benchmark highlighted that Diff- Dock struggles to predict binding poses for out of distribution binding pockets.^30^ To address this, we condition Eq. 1 on a putative binding site by biasing the translational component of the diffusion with the distance between the ligand’s center of geometry **r**, and the center of the binding site **r**_0_:

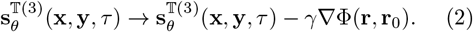

A flat-bottomed harmonic potential Φ characterizes the geometry and location of the binding site:

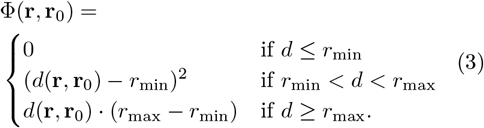

The function *d* measures the distance between **r** and **r**_0_, *r*_min_ is the inner radius, and the outer radius *r*_max_ limits the gradient for numerical stability. The strength of the potential is controlled by *γ*. When *γ* = 0 the generative process in Eq. 1, is *unconditional*, i.e. the center of geometry **r** freely explores the entire protein surface; in contrast, when *γ >* 0, the generative process is *conditional* and the center of geometry is biased by Φ towards a spherical region of radius *r*_min_ centered around **r**_0_. The orthogonality of the groups factoring ℙ guarantees that the rotational degrees of freedom of the generative process are unaffected by the translational bias.

### B. DiffDock–Glide

Since researchers are often interested in a small subset of representative poses, the samples from Diff- Dock ought to be scored and rank-ordered based on their plausibility. The original DiffDock implementation trains a separate neural network, called a confidence model, to predict a score that indicates if a pose is likely to be less than 2Å in root mean square deviation (RMSD) from a crystal pose. We take a different approach in using the ranking mechanism to introduce further inductive biases and evaluate pose quality without reference to the generative model or its training dataset.

Buttenschoen et. al. have previously shown that subtle features of real binding poses that are absent from the generated poses can be recovered by minimization under a molecular mechanics force field.^15^ Accordingly, we use Glide’s post-docking minimization pipeline to score and refine the generated poses. The pipeline optimizes ligand-receptor and intramolecular ligand interactions through rigid-body roto-translations and rotatable bonds optimizations under the OPLS-AA force field.^7^ The minimized complexes are then ranked using Glide’s scoring function, which has been empirically parameterized to approximate the binding free energy by rewarding or penalizing interactions, e.g. hydrogen bonding, van der Waals, hydrophobic, electrostatic interactions and internal strain energy.^7^

## III. APPLICATIONS

We explore two applications of the hybrid DiffDock– Glide model: pose prediction and virtual screening.

- Section III A evaluates DiffDock-Glide’s molecular docking performance on the Posebusters dataset. Our results indicate that binding site conditioning and energy minimization improve generalization to receptors outside the training set and increase the fraction of targets for which a pose less than 2Å in RMSD from the crystal pose is sampled. We also demonstrate a potential use case by using the *γ* = 1 model to dock against an allosteric binding site that the *γ* = 0 model does not recognize.
- In Section III B we use DiffDock-Glide to virtual screen binding and non-binding compounds from the DUD-E library against receptor structures generated by AF2. We report enrichments surpassing those obtained from DiffDock or Glide alone.

### A. Molecular docking

Molecular docking aims to predict the binding pose of a small molecule with a receptor. The task poses a significant challenge due to the diversity of physical and chemical interactions that contribute to binding stability. We benchmark DiffDock-Glide on the Posebusters molecular docking dataset.^15^

#### 1. Posebusters benchmark

The Posebusters (PB) dataset is a collection of protein-ligand complexes that is curated to serve as a benchmark for deep learning docking methods: PDBBind General Set v2020.^15^ The dataset provides 308 diverse, high-resolution complexes that are not present in PDBBind v2020. Besides the curated set of complexes, Posebusters implements consistency checks that valid binding poses must satisfy. These checks consider factors such as steric clashes, internal energy, and chemical consistency. A pose that passes all these checks is classified as PB-valid. Similarly, we define a Glide-valid (G-valid) condition by determining whether a pose meets Glide’s internal consistency requirements. Since Glide rejects poses with extreme values in the terms of its scoring function, any pose that receives a score from Glide is considered G-valid.

To compare the performance of different ranking methods we sample 40 poses for each PB target using DiffDock with *γ* = 0 and *γ* = 1. The poses are then ranked by the confidence model and Glide score after energy minimization. For the *γ* = 1 models, residues within 5Å of the crystallized ligand are considered part of the binding site, and their coordinates parameterize the binding site in Eq. 3: **r**_0_ is the center of geometry, *r*_min_ is three times the standard deviation of the pairwise inter-residue distances, and *r*_max_ is *r*_min_ + 10Å.

In Figure 1 we summarize the performance of DiffDock-Confidence (blue), DiffDock-Glide (red), and Glide Confgen (green) on the PB benchmark. The solid bars display the percentage of PB targets for which a crystal-like (<2Å RMSD) structure is the top ranked pose by each method, while the hatched bars denote the lowest RMSD selection among all sampled poses. The dashed green line denotes the performance of Glide Confgen, indicating that both DiffDock-Glide and DiffDock-Confidence fall short of state-of-the-art.

**FIG. 1.**
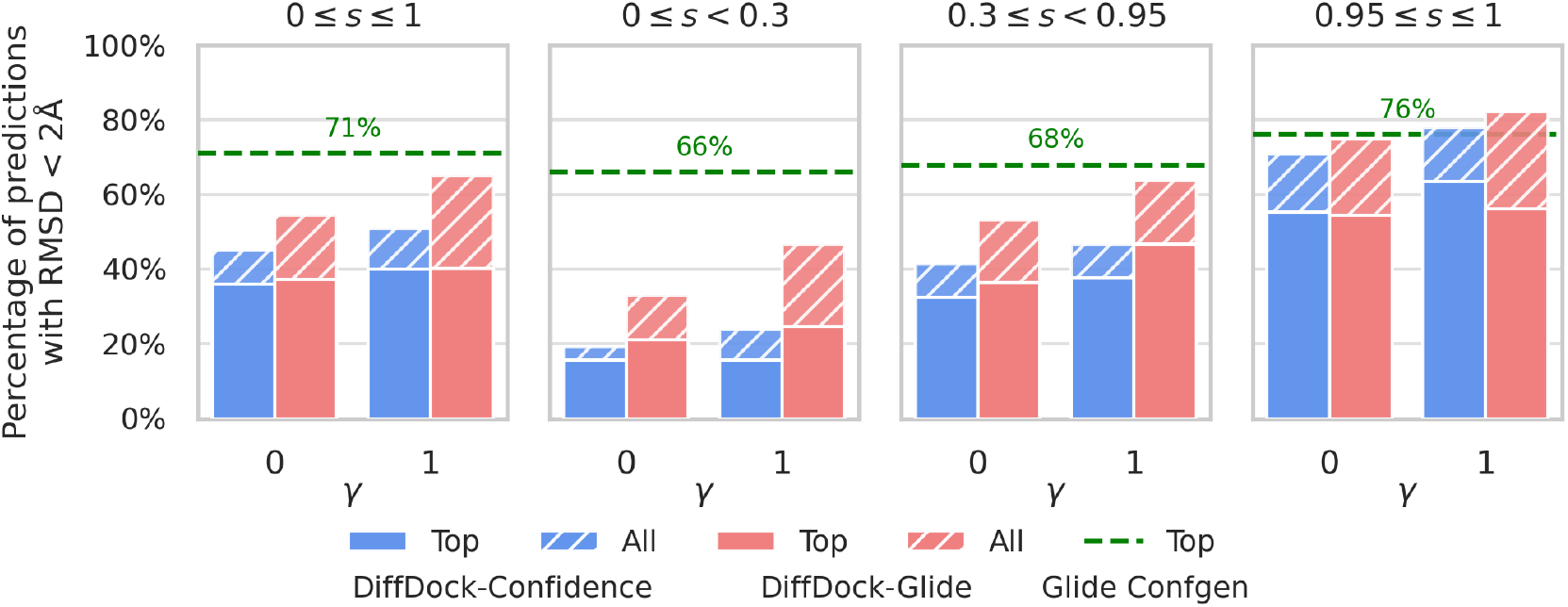
Performance of DiffDock-Glide on the Posebusters dataset. The percentage of Posebusters (PB) targets that have an RMSD within 2Å of the crystal pose is presented for three docking methods: DiffDock-Confidence, DiffDock-Glide, and Glide Confgen. For the DiffDock-based methods samples are generated with *γ* = 0 (no conditioning) and *γ* = 1 (binding site conditioning). The percentage of targets for which the top-ranked pose by the confidence model and Glide score (solid bars) is crystal-like is compared to the lowest RMSD selection among all poses (hatched bars). Glide Confgen (green line) is presented as a baseline. The leftmost panel summarizes performance for the entire PB dataset, and the remaining panels stratify the PB dataset by sequence similarity (*s*) to PDBBind General Set v2020.

We observe that *γ* = 1 has the expected effect of yielding a greater fraction of native-like poses compared to *γ* = 0. Likewise, we find that energy minimization increases the fraction of targets for which a crystal-like pose is sampled. Consistent with previous studies that identified artifacts in samples from DiffDock, we find that less than 0.1% of the sampled poses are G-valid without minimization.^15,16^ The most striking result is that the fraction of native-like poses selected according to the Glide score (solid red bars) is far below the ideal value (hatched red bars), reflecting the limitations of pose refinement via energy minimization.

In the remaining panels we stratify the results by maximum sequence similarity, *s*, to proteins in PDB- Bind v2020. We find that higher sequence similarity correlates with more accurate predictions, and the positive effect of energy minimization and binding site conditioning is greatest when *s <* 0.95. Notably, both DiffDock-Glide and DiffDock-Confidence are capable of sampling sub-2Å poses for a similar fraction of targets compared to Glide Confgen, provided that they have high (*s ≥* 0.95) sequence similarity.

#### 2. Docking into Allosteric Sites

Traditional small-molecule drug design operates on targeting the primary orthosteric binding site of a receptor to modulate its activity. Orthosteric sites typically exhibit well-defined, highly conserved pockets that bind endogenous signaling molecules with high specificity and affinity.^31^ However, certain “undruggable” proteins lack well-defined binding pockets, posing a significant challenge for traditional drug discovery. Allosteric modulation offers a compelling alternative by targeting allosteric sites that are spatially separate from the orthosteric site, but can nonetheless affect the protein’s function upon ligand binding.^32^ Targeting allosteric sites in G-protein coupled receptors, for instance, has led to improved target selectivity and reduced off-target effects.^33^

Although allosteric sites have therapeutic potential, current deep learning approaches remain focused on orthosteric targets. DiffDock’s training set contains both ortho- and allosteric complexes, but is dominated by the former due to the historical bias towards orthosteric drug discovery and limitations of structure probing techniques like X-ray crystallography, which readily resolve well-defined orthosteric sites but struggle to capture the dynamic and flexible structures of allosteric sites.^34^

In this section, we demonstrate how biasing Diff- Dock’s generative process enables targeting of allosteric sites. We design a cross-docking experiment centered around a *β*-2 Adrenergic G-coupled protein receptor domain containing both ortho- and allosteric sites. The reference complex (PDB: 6N48) is a *holo* crystal structure of the *β*-2 Adrenergic receptor in complex with two ligands: BI-167107 is bound to the orthosteric site while Compound-6FA (KBY) occupies the allosteric site, which is shallow and lies exposed on the membrane interface of the receptor.^35^ The ligand of interest, KBY, engages in *π*–*π* stacking interactions with side chain residue PHE1133 and interacts with hydrophobic residues on the membrane-exposed face of the receptor’s helical bundle. The target (PDB: 6KR8) is an NMR-derived model containing an ensemble of 10 conformers that represent the *apo* state.^36^ We select the first conformer as a representative *apo* structure, and backbone alignment with the *holo* conformer reveals an RMSD of less than 2Å. It is worth noting that the *holo* structure is present in the DiffDock training set whereas the *apo* structure is not, though it is included in 2019 version of the PDB.

Forty poses of KBY docked to the *apo* conformer were sampled using DiffDock-Confidence (*γ* = 0) and DiffDock-Glide (*γ* = 1) with the biasing potential centered on the allosteric site. Figure 2a depicts the distribution of poses sampled by both methods. The orange sphere sits at the centroid of the poses sampled by DiffDock-Confidence, all of which occupy the orthosteric site, while the blue sphere is the centroid of the poses generated by DiffDock-Glide, all of which are within 5Å of the allosteric binding site after aligning the *holo* ligand (yellow sphere). In this example, the unguided DiffDock-Confidence model always docks KBY at the orthosteric site. The introduction of a simple biasing potential to the generative process in DiffDock-Glide enables one to target the allosteric site.

**FIG. 2.**
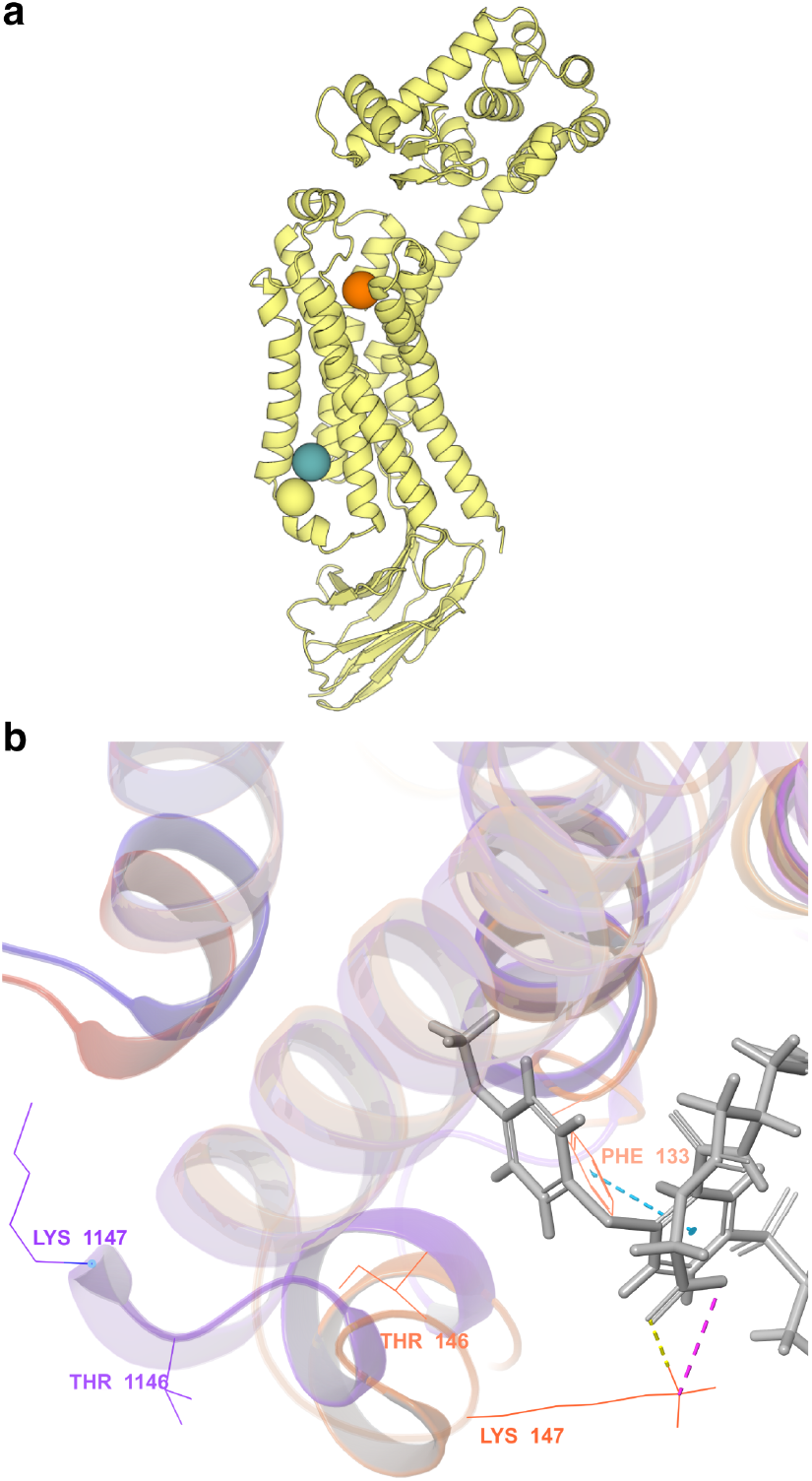
Targeting the allosteric site of *β*-2 Adrenergic receptor with ligand KBY. (a) The *holo* structure (PDB: 6N48) is shown. DiffDock-Confidence (*γ* = 0) samples all 40 generated poses at the orthosteric site (orange sphere), while DiffDock-Glide (*γ* = 1) samples poses (denoted by the blue sphere) within 5Å of the *holo* allosteric site (yellow sphere). (b) The allosteric sites of the *holo* (orange) and *apo* (purple) receptors are aligned and su- perimposed alongside a pose sampled by DiffDock-Glide (*γ* = 1). Stabilizing interactions recovered by Glide are indicated by dotted lines.

Figure 2b depicts a pose sampled by DiffDock-Glide (*γ* = 1) alongside the aligned *holo* (orange) and *apo* (purple) allosteric sites. The *apo* intracellular loop 2 (THR146 and LYS147) is in a different orientation relative to the *holo* conformation, resulting in a narrower binding site. Consequently, we expect the final refined pose to deviate from the *holo* ligand. Indeed, Glide resolves clashes with THR146 and LYS147 during post-docking minimization by including protein side chain residues that are not present in DiffDock, and, upon aligning the *holo* binding site to the *apo* conformer, we find that the DiffDock-Glide pose deviates from the *holo* pose. Still, a primary interaction is shared between the *holo* and DiffDock-Glide poses: both form *π*-*π* stacking with residue PHE133 (blue dotted line). The DiffDock-Glide pose forms additional stabilizing interactions between the ligand and LYS147: a hydrogen bond (yellow dotted line) and a salt bridge (magenta dotted line) form between the ligand’s carboxylate and the positively charged *ϵ*-amino group on the side chain residue.

### B. Virtual screening

*In silico* hit identification uses computational methods to discover potential drug candidates, or hits, from chemical libraries. A successful virtual screening effort effectively prioritizes binding ligands (actives) over non-binders (decoys). Two classes of techniques are often used to screen large libraries for compounds likely to bind to a target protein.

Ligand-based methods are typically used for lead compound optimization when the structure of the target protein is unknown. They rely on chemical features of known actives to search for compounds with similar binding modalities. Techniques like quantitative structure-activity relationship (QSAR) are particularly effective in prioritizing biologically active compounds by using properties of other known actives.^37^

On the other hand, structure-based approaches like molecular docking rely on high-resolution protein structures to predict the binding interactions of compounds. Classical docking tools explore potential binding modes by optimizing the orientation of the candidate ligand in the protein’s binding site according to a scoring function. Well-known tools like AutoDock Vina^5,38^, DOCK^8^, Glide^7^, and GOLD^6^ use distinct algorithms for both pose generation and scoring.^39^ In the following section, we explore if DiffDock-Glide is suitable for structure-based virtual screening.

#### 1. AF2-DUD-E benchmark

Most molecular docking tools perform best on high-quality ligand-receptor structures when searching for binding poses. DiffDock, however, is unique in that it infers binding poses solely based on the *C*_*α*_ co- ordinates. Accordingly, we believe that DiffDock-Glide should perform well when screening against low-quality receptor structures, e.g. those with mis- placed sidechains. To test our hypothesis, inspired by a previous study that showed AF2-generated receptor structures are not generally suitable for virtual screening with molecular docking^40^, we screen active and decoy compounds in the DUD-E dataset against receptor structures generated by AF2 using DiffDock- Confidence, DiffDock-Glide, and Glide Confgen.

The DUD-E dataset is a standard benchmark in virtual screening for evaluating a method’s ability to distinguish between actives and decoy compounds.^28^ It contains proteins from chemically diverse families, each with a set of validated active compounds and physicochemically similar but structurally distinct decoy compounds.

We screen against the 27 structures from the AF2 database for which the corresponding DUD-E structure does not contain cofactors at the binding site or clashes with the *holo* ligand. The structures are prepared using the truncation procedure outlined by Zhang et. al.^40^ For each structure-compound pair, five poses are sampled, minimized, and scored with the same parameters as the docking study in Section III A, except that residues constituting the binding site are chosen based on the DUD-E crystal structure and *r*_min_ and **r**_0_ are derived from the corresponding residue geometry in the AF2 structure.

Table I reports the average EF^41^ and BEDROC^42^ (*α* = 160.9) enrichments. Intuitively, EF1% measures how concentrated actives are in the top 1% of rankings compared to the entire set of compounds. Similarly, BEDROC (*α* = 160.9) non-linearly emphasizes early enrichment due to actives among the highest ranked compounds. For both measures, a larger value indicates a greater concentration of active compounds among the top ranks. The disproportionately low enrichment values for DiffDock-Confidence (*γ* = 0) are due to it being a blind screening method. The corresponding DiffDock-Glide (*γ* = 0) model is not blind since Glide penalizes the rank of poses generated outside the binding site. The top performing DiffDock-Glide (*γ* = 1) model has the most inductive biases among all the DiffDock-based methods, and even outperforms Glide Confgen. Figure 3 compares enrichment on a per-target basis for Glide Confgen, DiffDock-Glide (*γ* = 1), and DiffDock-Confidence (*γ* = 1). While the maximum observed enrichment across all targets from Glide Confgen, DiffDock-Glide (*γ* = 1) provides the greatest enrichment for over half of the targets. It is also noteworthy that the purely data-driven DiffDock-Confidence (*γ* = 1) model out- performs Glide Confgen for seven targets.

**TABLE I.** Average enrichments from virtual screening DUD-E compounds against AF2-generated receptors. The EF values are computed with a top 1% cutoff. Like- wise, the BEDROC values are computed with *α* = 160.9 to emphasize enrichment in the top 1% of rankings.

| Screening Method | EF (↑) | BEDROC (↑) |
| --- | --- | --- |
| DiffDock-Confidence ($\gamma = 0$) | 1.07 | 0.02 |
| DiffDock-Confidence ($\gamma = 1$) | 9.24 | 0.16 |
| Glide Confgen | 13.1 | 0.23 |
| DiffDock-Glide ($\gamma = 0$) | 15.5 | 0.28 |
| DiffDock-Glide ($\gamma = 1$) | **16.2** | **0.31** |

**FIG. 3.**
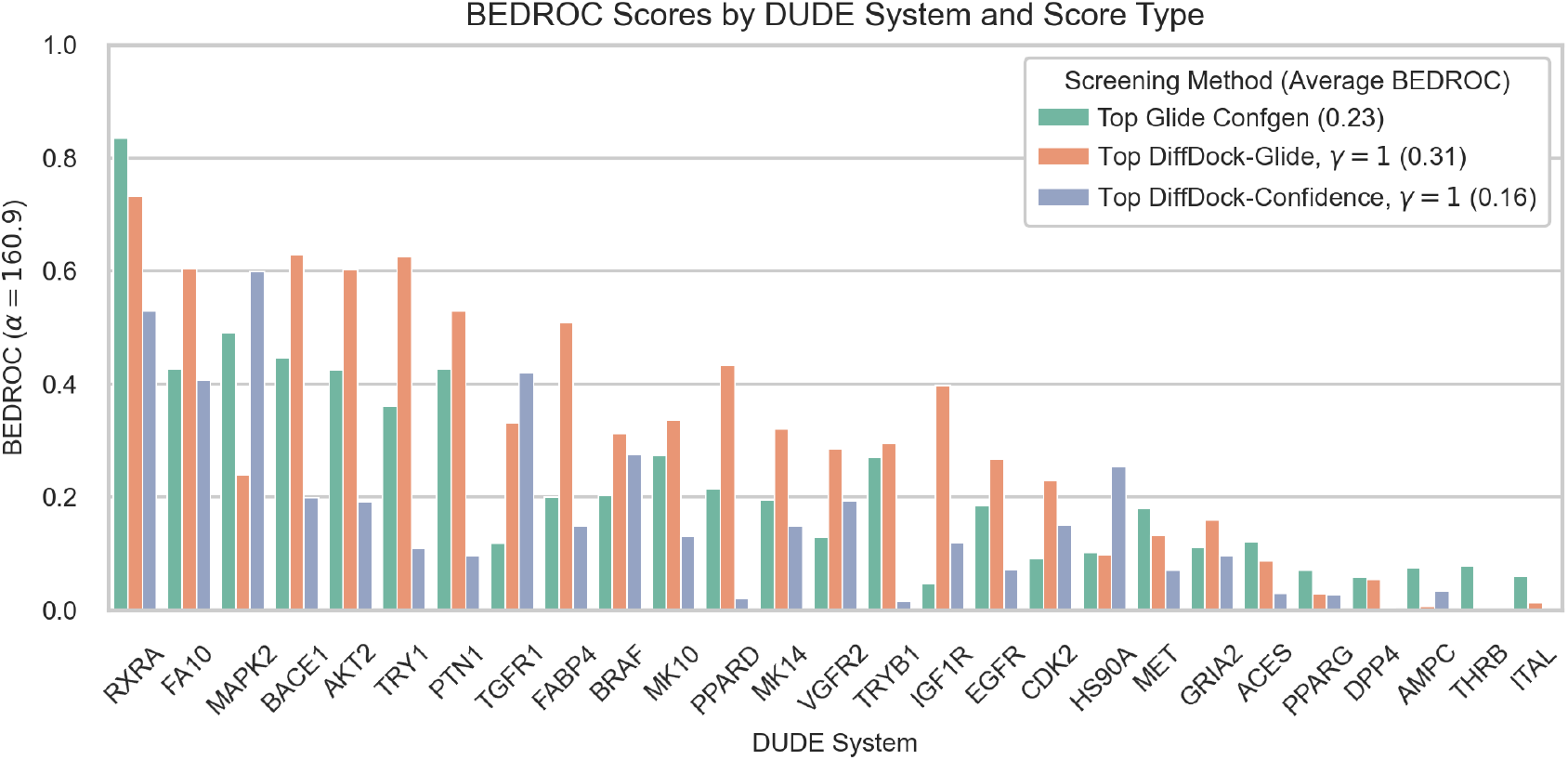
Target-specific enrichment for the AF2-DUD-E virtual screening benchmark. BEDROC (*α* = 160.9) enrichments are displayed for the 27 AF2-DUD-E target. Five poses are sampled per compound by DiffDock, and scored by the confidence model and Glide’s post-docking minimization pipeline. Compounds are ranked according to the top confidence (blue) and Glide (orange) score among the five poses for the BEDROC calculation. BEDROC enrichments from Glide Confgen (green) are provided as a reference.

#### 2. Case Study for IGF-1R

DiffDock-Glide demonstrates significantly higher enrichment of active compounds compared to Glide Confgen for the IGF-1R virtual screening target (see Figure 3). Glide scores of the top 1% of ranked compounds range from − 9.5 to − 7.6 for Glide Confgen and − 9.1 to − 6.9 for DiffDock-Glide (*γ* = 1), indicating that both methods sample favorable binding poses. For reference, scores greater than 6 are generally associated with poor docking poses.^7^

Although the ranges of scores are similar for both methods, the chemical composition of their top 1% compounds differs. DiffDock-Glide ranks 39 active compounds with 33 unique scaffolds among the top 1%. Notably, 91.4% of the top 1% compounds contain pyrimidine-pyrrole cores. Glide Confgen, on the other hand, recovers 11 actives with 11 unique scaf- folds in the top 1% and does not show preference for any particular DUD-E chemical series: only two of the top-ranked actives have pyrimidine-pyrrole cores, and the others include, for example, thiazoles and benzimidazoles. Even though the composition of the top 1% of compounds is different from DiffDock-Glide, Glide Confgen still assigns valid (i.e. negative) scores to all the top actives identified by DiffDock-Glide, which suggests that the Glide score is unable to prioritize actives over decoys in this instance. A more detailed analysis reveals that Glide Confgen fails to model the formation of buried hydrogen bonds with nitrogen acceptors. The scores for IGF-1R are dominated by hydrophobic contributions that are not balanced by suitable penalties for protein desolvation. This is a known limitation of Glide Confgen and is addressed by GlideWS, formerly known as WScore.^9^

To demonstrate the improvement that pose minimization brings to DiffDock-Glide, we analyze performance for active compound CHEMBL461876 with and without minimization. Figure 4a depicts the un- minimized (gray) and minimized (purple) poses. The un-minimized pose undermines interpretation of the binding mode due to an over-packed binding pocket and unrealistic contacts with residues ARG1003 and LEU1005. Atom C17 of the ligand clashes with the guanidinium group of ARG1003 and CD2 of LEU1005 due to interatomic distances less than 3Å. Pose minimization improves the rank of CHEMBL461876 from 372 to 3 by resolving the steric clashes.

**FIG. 4.**
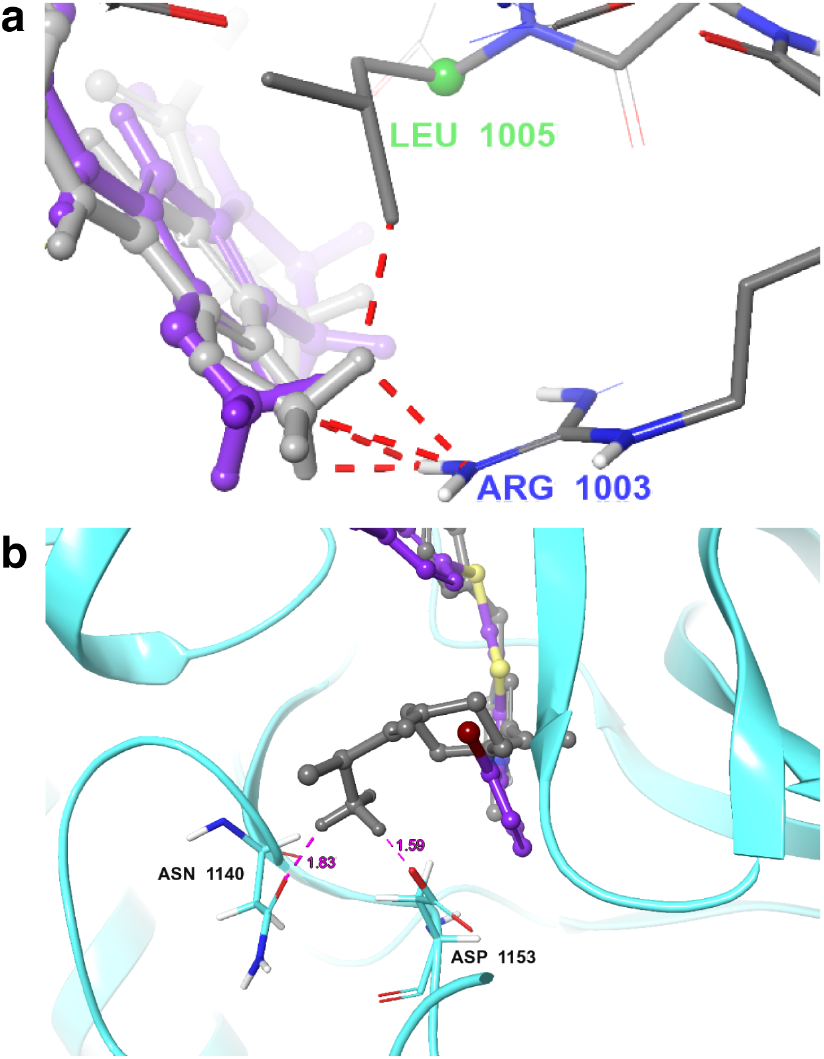
Analysis of poses sampled for IGF-1R. (a) DiffDock-Glide poses before (gray) and after (pur- ple) minimization are shown for ligand CHEMBL461876. Interatomic distances less than 3Å (red dotted lines) in- dicate clashes between the receptor and DiffDock pose at LEU1005 and ARG1003. (b) The DiffDock-Glide pose (purple) for CHEMBL251795 fails to establish a potent bidentate salt bridge with residues ASN1140 and ASP1153 in the active site. The Glide-Confgen pose (gray) is correctly oriented to form the salt-bridge. Inter- actions with interatomic distances measuring less than 3.5Å are depicted as magenta dotted lines.

Still, there are instances where Glide Confgen successfully models essential interactions that DiffDock-Glide fails to capture. We focus on active compound CHEMBL251795, which is stabilized by a buried salt-bridge formed between the protonated amide and charged carboxylate on residue ASP1153, and a hydrogen bond between the ligand and ASN1140. Figure 4b depicts poses sampled by Glide Confgen (gray) with the bidentate salt-bridge and DiffDock-Glide (purple) without this key interaction. The top Glide Confgen pose is ranked as the top active with a score of −9.5. DiffDock-Glide does not sample the salt-bridge, and although minimization proceeds successfully, it yields a comparatively poor score of−4.7.

## IV. OUTLOOK

In this work we have developed a hybrid approach to molecular docking called DiffDock-Glide that improves the out-of-distribution performance of data-driven docking. The core idea is to use a deep generative model to sample from a learned distribution of binding poses and subsequently relax the sampled poses with a physics and chemistry-based energy function that contains inductive biases. We conclusively demonstrate that providing such biases dramatically improves the performance of DiffDock. The primary limitations of DiffDock-Glide are sampling crystal-like poses, especially for out-of-distribution receptors, and selecting a crystal-like pose from the sampled distribution.

We found that the confidence and Glide scores are unable to reliably select for such poses even if they are present among the generate samples. Physics-based pose refinement beyond energy minimization may lead to more accurate pose ranking. We also found that DiffDock-Glide is especially suitable for virtual screening against AF2 receptor structures, though it should be noted that all the screened receptors except *PPARD* are found in DiffDock’s training set. At the time of writing, there is no virtual screening equivalent of the Posebusters set, so the effects of overfitting could not be systematically investigated. Though based on the Posebusters analysis, we anticipate that DiffDock’s performance will degrade for out-of-distribution targets, and the decline can be mitigated via the inclusion of appropriate biases.

Co-folding models that predict the structure of entire complexes have garnered much attention since the publication of DiffDock. Nevertheless, we still believe DiffDock to be the most suitable base model for our approach: while models like Boltz1^43^ and AlphaFold3^12^ are indeed diffusion models, their dynamics unfold in the full coordinate space and are less straightforward to control compared to DiffDock’s torsional representation which respects molecular symmetries.

On a final note, we would like to emphasize that our modifications to DiffDock’s generative dynamics are independent of the model’s weights and therefore do not require retraining to implement, nor do they present substantial computational overhead during sampling. Overall, we introduce DiffDock-Glide as a practical solution for structure-based drug discovery applications that incorporates well-validated biophysical principles, maintains the advantages of learned representations, and is more interpretable compared to fully deep learning approaches.

## V. SOFTWARE AND DATA

All Glide calculations were performed using Schrödinger Suite (2023-2 release). The protein protonation state is assigned with PROPKA33,34 at pH 7.4. DUD-E actives and decoys were processed using the LigPrep program of Schrödinger Suite starting from a SMILES representation of each molecule, using the following settings: (1) Tautomers were generated using Schrödinger’s Epik with a target pH of 7.0 +/- 1.0 and (2) a maximum of 32 stereoisomers were generated for each molecule. The Posebusters (https://zenodo.org/records/8278563) dataset was used for molecular docking, and the DUD-E (http://dude.docking.org) dataset and AF2 database (https://alphafold.ebi.ac.uk) for virtual screening.

## References

1 A. C. Anderson, “The process of structure-based drug design,” Chemistry & biology 10, 787–797 (2003).

2 B. K. Shoichet, “Virtual screening of chemical libraries,” Nature 432, 862–865 (2004).

3 N. S. Pagadala, K. Syed, and J. Tuszynski, “Software for molecular docking: a review,” Biophysical reviews 9, 91–102 (2017).

4 K. Klarich, B. Goldman, T. Kramer, P. Riley, and W. P. Walters, “Thompson sampling an efficient method for searching ultralarge synthesis on demand databases,” Journal of Chemical Information and Modeling 64, 1158–1171 (2024).

5 O. Trott and A. J. Olson, “Autodock vina: improving the speed and accuracy of docking with a new scoring function, efficient optimization, and multithreading,” Journal of computational chemistry 31, 455–461 (2010).

6 M. L. Verdonk, J. C. Cole, M. J. Hartshorn, C. W. Murray, and R. D. Taylor, “Improved protein–ligand docking using gold,” Proteins: Structure, Function, and Bioinformatics 52, 609–623 (2003).

7 R. A. Friesner, J. L. Banks, R. B. Murphy, T. A. Halgren, J. J. Klicic, D. T. Mainz, M. P. Repasky, E. H. Knoll, M. Shelley, J. K. Perry, et al., “Glide: a new approach for rapid, accurate docking and scoring. 1. method and assessment of docking accuracy,” Journal of medicinal chemistry 47, 1739–1749 (2004).

8 W. J. Allen, T. E. Balius, S. Mukherjee, S. R. Brozell, D. T. Moustakas, P. T. Lang, D. A. Case, I. D. Kuntz, and R. C. Rizzo, “Dock 6: Impact of new features and current docking performance,” Journal of computational chemistry 36, 1132–1156 (2015).

9 R. B. Murphy, M. P. Repasky, J. R. Greenwood, I. Tubert-Brohman, S. Jerome, R. Annabhimoju, N. A. Boyles, C. D. Schmitz, R. Abel, R. Farid, et al., “Wscore: a flexible and accurate treatment of explicit water molecules in ligand–receptor docking,” Journal of medicinal chemistry 59, 4364–4384 (2016).

10 J. Jumper, R. Evans, A. Pritzel, T. Green, M. Figurnov, O. Ronneberger, K. Tunyasuvunakool, R. Bates, A. Žídek, A. Potapenko, et al., “Highly accurate protein structure prediction with alphafold,” nature 596, 583–589 (2021).

11 R. Krishna, J. Wang, W. Ahern, P. Sturmfels, P. Venkatesh, I. Kalvet, G. R. Lee, F. S. Morey-Burrows, I. Anishchenko, I. R. Humphreys, et al., “Generalized biomolecular modeling and design with rosettafold all-atom,” Science 384, eadl2528 (2024).

12 J. Abramson, J. Adler, J. Dunger, R. Evans, T. Green, A. Pritzel, O. Ronneberger, L. Willmore, A. J. Ballard, J. Bambrick, et al., “Accurate structure prediction of biomolecular interactions with alphafold 3,” Nature, 1–3 (2024).

13 G. Corso, H. Stärk, B. Jing, R. Barzilay, and T. Jaakkola, “Diffdock: Diffusion steps, twists, and turns for molecular docking,” arXiv preprint 2210.01776 (2022).

14 Z. Qiao, W. Nie, A. Vahdat, T. F. Miller III, and A. Anandkumar, “State-specific protein–ligand complex structure prediction with a multiscale deep generative model,” Nature Machine Intelligence 6, 195–208 (2024).

15 M. Buttenschoen, G. M. Morris, and C. M. Deane, “Pose-busters: Ai-based docking methods fail to generate physically valid poses or generalise to novel sequences,” Chemical Science 15, 3130–3139 (2024).

16 D. Errington, C. Schneider, C. Bouysset, and F. A. Dreyer, “Assessing interaction recovery of predicted protein-ligand poses,” arXiv preprint 2409.20227 (2024).

17 S. Geman, E. Bienenstock, and R. Doursat, “Neural networks and the bias/variance dilemma,” Neural computation 4, 1–58 (1992).

18 T. Yoon, J. Y. Choi, S. Kwon, E. K. Ryu, “Diffusion probabilistic models generalize when they fail to memorize,” in ICML 2023 Workshop on Structured Probabilistic Inference (2023) and Generative Modeling, 2023.

19 T. M. Mitchell, “The need for biases in learning generalizations,” (1980).

20 F. Fuchs, D. Worrall, V. Fischer, and M. Welling, “Se (3)-transformers: 3d roto-translation equivariant attention networks,” Advances in neural information processing systems 33, 1970–1981 (2020).

21 A. Musaelian, S. Batzner, A. Johansson, L. Sun, C. J. Owen, M. Kornbluth, and B. Kozinsky, “Learning local equivariant representations for large-scale atomistic dynamics,” Nature Communications 14, 579 (2023).

22 E. Hoogeboom, V. G. Satorras, C. Vignac, and M. Welling, “Equivariant diffusion for molecule generation in 3d,” in International conference on machine learning (PMLR, 2022) pp. 8867–8887.

23 B. Jing, G. Corso, J. Chang, R. Barzilay, and T. Jaakkola, “Torsional diffusion for molecular conformer generation,” Advances in Neural Information Processing Systems 35, 24240–24253 (2022).

24 V. De Bortoli, E. Mathieu, M. Hutchinson, J. Thornton, Y. W. Teh, and A. Doucet, “Riemannian score-based generative modelling,” Advances in Neural Information Processing Systems 35, 2406–2422 (2022).

25 G. Corso, Y. Xu, V. De Bortoli, R. Barzilay, and T. Jaakkola, “Particle guidance: non-iid diverse sampling with diffusion models,” arXiv preprint 2310.13102 (2023).

26 L. Herron, K. Mondal, J. S. Schneekloth Jr, and P. Tiwary, “Inferring phase transitions and critical exponents from limited observations with thermodynamic maps,” Proceedings of the National Academy of Sciences 121, e2321971121 (2024).

27 S. Lee, R. Wang, L. Herron, and P. Tiwary, “Exponentially tilted thermodynamic maps (exptm): Predicting phase transitions across temperature, pressure, and chemical potential,” (2025).

28 M. M. Mysinger, M. Carchia, J. J. Irwin, and B. K. Shoichet, “Directory of useful decoys, enhanced (dud-e): better ligands and decoys for better benchmarking,” Journal of medicinal chemistry 55, 6582–6594 (2012).

29 Y. Song, J. Sohl-Dickstein, D. P. Kingma, A. Kumar, S. Ermon, and B. Poole, “Score-based generative modeling through stochastic differential equations,” arXiv preprint 2011.13456 (2020).

30 G. Corso, A. Deng, B. Fry, N. Polizza, R. Brazilay, and T. Jaakkola, “Deep confident steps to new pockets: Strategies for docking generalization,” International Conference on Learning Representations (2024).

31 R. Nussinov and C.-J. Tsai, “The different ways through which specificity works in orthosteric and allosteric drugs,” Current pharmaceutical design 18, 1311–1316 (2012).

32 X. Xie, T. Yu, X. Li, N. Zhang, L. J. Foster, C. Peng, W. Huang, and G. He, “Recent advances in targeting the “undruggable” proteins: from drug discovery to clinical trials,” Signal transduction and targeted therapy 8, 335 (2023).

33 R. Nussinov, M. Zhang, Y. Liu, and H. Jang, “Alphafold, allosteric, and orthosteric drug discovery: Ways forward,” Drug discovery today 28, 103551 (2023).

34 D. S. Vora and S. Yadav, “Fantastic allosteric binding sites and why deep learning cannot find them,” in I Can’t Believe It’s Not Better: Challenges in Applied Deep Learning (2025).

35 X. Liu, A. Masoudi, A. W. Kahsai, L.-Y. Huang, B. Pani, D. P. Staus, P. J. Shim, K. Hirata, R. K. Simhal, A. M. Schwalb, et al., “Mechanism of β2ar regulation by an intracellular positive allosteric modulator,” Science 364, 1283–1287 (2019).

36 S. Imai, T. Yokomizo, Y. Kofuku, Y. Shiraishi, T. Ueda, and I. Shimada, “Structural equilibrium underlying ligand-dependent activation of β2-adrenoreceptor,” Nature Chemical Biology 16, 430–439 (2020).

37 M. Butkiewicz, E. W. Lowe, R. Mueller, J. L. Mendenhall, P. L. Teixeira, C. D. Weaver, and J. Meiler, “Benchmarking ligand-based virtual high-throughput screening with the pubchem database,” Molecules 18, 735–756 (2013).

38 J. Eberhardt, D. Santos-Martins, A. F. Tillack, and S. Forli, “Autodock vina 1.2. 0: New docking methods, expanded force field, and python bindings,” Journal of chemical information and modeling 61, 3891–3898 (2021).

39 A. C. O. O. Agu, P.C., “Molecular docking as a tool for the discovery of molecular targets of nutraceuticals in diseases management,” Sci Rep 13 (2023), 10.1038/s41598-023-40160-2.

40 Y. Zhang, M. Vass, D. Shi, E. Abualrous, J. M. Chambers, N. Chopra, C. Higgs, K. Kasavajhala, H. Li, P. Nandekar, et al., “Benchmarking refined and unrefined alphafold2 structures for hit discovery,” Journal of Chemical Information and Modeling 63, 1656–1667 (2023).

41 D. A. Pearlman and P. S. Charifson, “Improved scoring of ligand- protein interactions using owfeg free energy grids,” Journal of medicinal chemistry 44, 502–511 (2001).

42 J.-F. Truchon and C. I. Bayly, “Evaluating virtual screening methods: good and bad metrics for the “early recognition” problem,” Journal of chemical information and modeling 47, 488–508 (2007).

43 J. Wohlwend, G. Corso, S. Passaro, M. Reveiz, K. Leidal, W. Swiderski, T. Portnoi, I. Chinn, J. Silterra, T. Jaakkola, and R. Barzilay, “Boltz-1: Democratizing biomolecular interaction modeling,” bioRxiv (2024), 10.1101/2024.11.19.624167.

